# The preeminent role of directional selection in generating extreme morphological change in Glyptodonts (Cingulata; Xenarthra)

**DOI:** 10.1101/2021.07.07.451530

**Authors:** Fabio A. Machado, Gabriel Marroig, Alex Hubbe

**Affiliations:** Department of Biology, Virginia Tech, USA; Departamento de Genética e Biologia Evolutiva, Instituto de Biociências, Universidade de São Paulo, Brazil; Departamento de Oceanografia, Instituto de Geociências, Universidade Federal da Bahia, Brazil

**Keywords:** Skull, Extreme morphology, Fossil, Natural selection, Rates of Evolution

## Abstract

The prevalence of stasis on macroevolution has been classically taken as evidence of the strong role of stabilizing selection in constraining morphological evolution. Rates of evolution calculated over longer time scales tend to fall below the expected under genetic drift, suggesting that the signal for directional selection is erased at longer time scales. Here we investigated the rates of morphological evolution of the skull in a fossil lineage that underwent extreme morphological modification, the glyptodonts. Contrary to what was expected, we show here that directional selection was the main process during the evolution of glyptodonts. Furthermore, the reconstruction of selection patterns shows that traits selected to generate a glyptodont morphology are markedly different from those operating on extant armadillos. Changes in both direction and magnitude of selection are probably tied to glyptodonts’ invasion of a specialist-herbivore adaptive zone. These results suggest that directional selection might have played a more important role in the evolution of extreme morphologies than previously imagined.

## Introduction

Identifying signals of adaptive evolution using rates of change on a macroevolutionary scale is inherently problematic because tempo and mode of evolution are intertwined. While stabilizing selection slows down phenotypic change, directional selection can produce faster rates of evolution (1– 3). This means that the action of selection (directional or stabilizing) can be identified by contrasting how fast species evolve to the expected rates under genetic drift (1, 2). While this framework has been successfully applied to microevolutionary time scales (4), its usefulness to macroevolutionary and deep-time studies has been contentious (5).

At larger time scales, it is challenging to quantify the effect of directional selection on the evolutionary process. The leading mechanism of phenotypic evolution on the macroevolutionary scale is thought to be stabilizing selection, as attested by the prevalence of stasis in the fossil record (6). This does not necessarily mean that directional selection did not play any role in phenotypic diversification. In fact, because the adaptive landscape (the relationship between phenotype and fitness) is thought to be rugged, with many different optimal phenotypic combinations (peaks), evolutionary change under directional selection is bound to be fast during peak shifts and punctuated at the macroevolutionary scale (Fig. 1). Thus, because stasis is so prevalent and measured rates of change reflect the net-evolutionary processes that operated during the course of lineages’ histories, rates of evolution calculated on phylogenies and fossil record tend to be small compared to rates observed in extant populations (5). As a result, measured net-evolutionary rates of morphological change are more commonly than not consistent with evolution under stabilizing selection or drift (3, 5, 7–10). This probably explains why macroevolutionary studies usually infer the action of directional selection indirectly through pattern-based methods (e.g., through peak-transitions in diffusion-like models (11); the comparison of rates of change between lineages (12), etc.), rather than adopting biologically informed models (e.g., those derived from quantitative genetics theory).

**Fig. 1.**
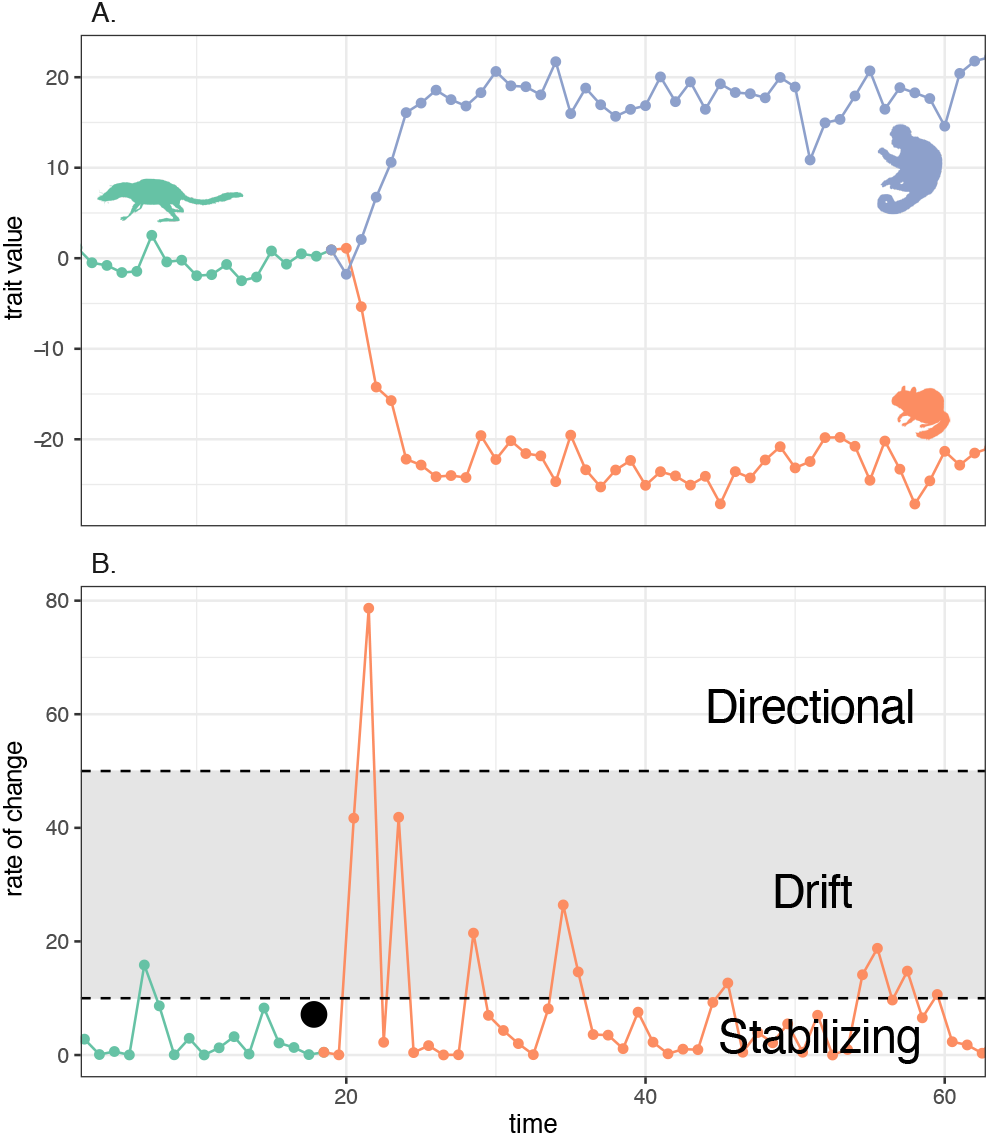
**A**. Evolution of a trait in an ancestral population (green) and during cladogenesis caused by a peak shift, resulting in two daughter lineages (blue and orange). **B**. Rates of evolution of the trait depicted in **A** for the ancestral species and one of the daughter lineages. The gray band represents the rates of evolution consistent with the expectation under genetic drift. Values above it are produced by directional selection and values below it by stabilizing selection. Even though the linage experienced expressive directional selection, in practice, one calculates net rates of evolution over the whole period since the origin of the species (black dot), which tend to be small and consistent with the expected under stabilizing selection.

While this departure from a biologically informed model might be justified for univariate traits (5, 7, 10), there is a wealth of evidence showing that the evolution of multivariate complex phenotypes, such as the mammalian skull, follow rules of growth and inheritance that can be modeled using quantitative genetics theory (9, 13, 14). Complex morphological traits are composed of subunits that are interconnected to each other to varying degrees. These different degrees of interconnection, generated by the complex interaction of multiple ontogenetic pathways, lead to an unequal variance distribution among parts (15), which in turn results in constraints to adaptive evolution (1, 3, 16, 17). Ignoring the possible role of constraints on the evolution of morphology can have many potential drawbacks, including the establishment of unrealistic expectations for drift in certain directions of the morphospace (3, 18) and possibly obscuring the signal of directional selection. To investigate the effect of directional selection on macroevolutionary scales, while avoiding potential pitfalls inherent to the analysis of multivariate traits, one should aim at studying a complex trait with a well-known trait covariance structure and that probably experienced high rates of evolutionary change. One such example is the case of the evolution of the glyptodont skull.

Glyptodonts originated in South America and are members of the order Cingulata (Xenarthra, Mammalia), which includes the extant armadillos. An emblematic characteristic of glyptodonts that is shared with all members of this order is the presence of an osteoderm-based exoskeleton (19). However, glyptodonts have a set of unique traits that makes them stand out from all other mammals, including defensive tail weaponry, large size, and, specifically, a highly modified skull. Glyptodonts skull has undergone a particular process of telescoping in which the rostrum is ventrally expanded, and the tooth row is posteriorly extended, resulting in a structure with unprecedented biomechanical proprieties among mammals (20, 21). Because of this unique morphology, glyptodonts’ phylogenetic position has been historically uncertain (19, 22, 23). Classically, they have been regarded as a sister group to extant armadillos. However, recent ancient DNA analysis support that glyptodonts are nested within the extant armadillo clade, having originated during the late Miocene (24, 25) (Fig. 2). As such, glyptodonts evolved during the same period of time it took for different extant armadillo lineages to diversify, making this a perfect model to study morphological evolution in a restricted taxonomic scope. Furthermore, the covariance structure of cranial traits has been extensively investigated in mammals in general and in Xenarthra specifically (26), allowing one to investigate the evolution of cranial morphology in Glyptodonts while considering the potential constraining effects of among-traits integration.

**Fig. 2.**
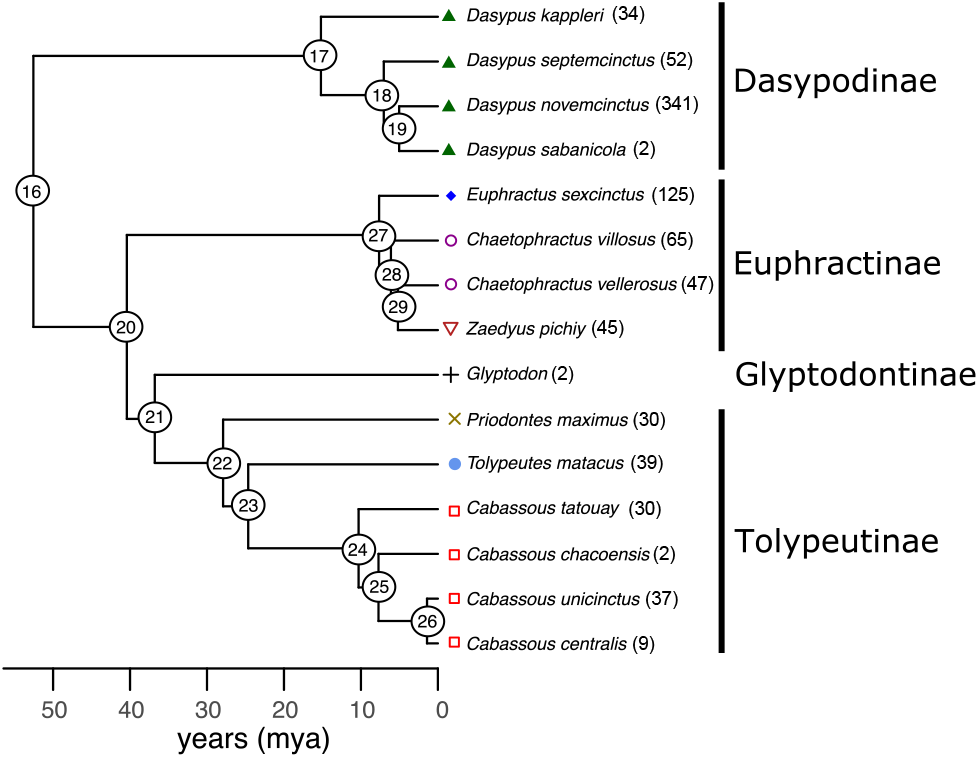
Cingulata phylogeny used in the present analysis, including the extinct *Glyptodon* based on (24). Numbers represent the node names referred throughout our analyses. Colors and symbols are using in plot results.

The present contribution aims to use the unique glyptodont case to evaluate if we can identify the signal of directional selection on multivariate macroevolutionary data using biologically informed models of morphological evolution. Specifically, we calculate the empirical rates of evolution among Cingulata taxa and compare that to the expected rates of evolution under genetic drift to identify cases of morphological evolution that can be confidently associated with directional selection. Additionally, we retrospectively estimate the selective forces necessary to generate all morphological diversification within Cingulata to understand better if evolutionary processes responsible for the evolution of glyptodonts are qualitatively different (i.e., selection guided glyptodonts in different directions) from those associated with other Cingulata taxa.

## Materials and Methods

### Morphometrics and trait covariance

Morphometric analysis was based on 33 linear distances obtained from 3D cranial landmarks (Fig. S1) digitized from 860 museum collection specimens (Table S1). These measurements are designed to capture general dimensions of bones and structures of the mammalian skull. Landmarks and distances are based on (26). Traits were log-transformed to normalize differences in variance due to scale differences. Sample sizes ranged from 2—341 per species (Table S1). We adopted the phenotypic covariance matrix (or **P**) as a surrogate of the additive genetic covariance matrix **G** (see (26) for details).

Here, we calculated **P** as the pooled within-group covariance matrix for all species. This assumes that the **P** is stable throughout the group’s evolution (26). Expressly, we assume that the **P** for glyptodonts is the same as for the rest of Cingulata. While this might be contentious due to the extreme morphological modification in the group, **P**s were shown to be similar among extant and fossil Xenarthra, including giant ground sloths (26). Additionally, because glyptodonts are nested within extant Cingulata, using the pooled covariance for living species as a model for glyptodont can be seen as a case of phylogenetic bracketing (27).

Form (shape+size) was quantified as the full morphospace, while shape was obtained by projecting out the leading PC of the within-group **P** the dataset. Given that PC1 is usually a size direction (i.e., where all loadings show similar values and the same sign 28; Tab. S2), this produces a dataset that is free from both size and allometric variation. Size was quantified as the geometric mean of all traits for each individual. Morphological variation for form was described with a principal component analysis (PCA) for cranial data (Fig 3, Tab S2).

**Fig. 3.**
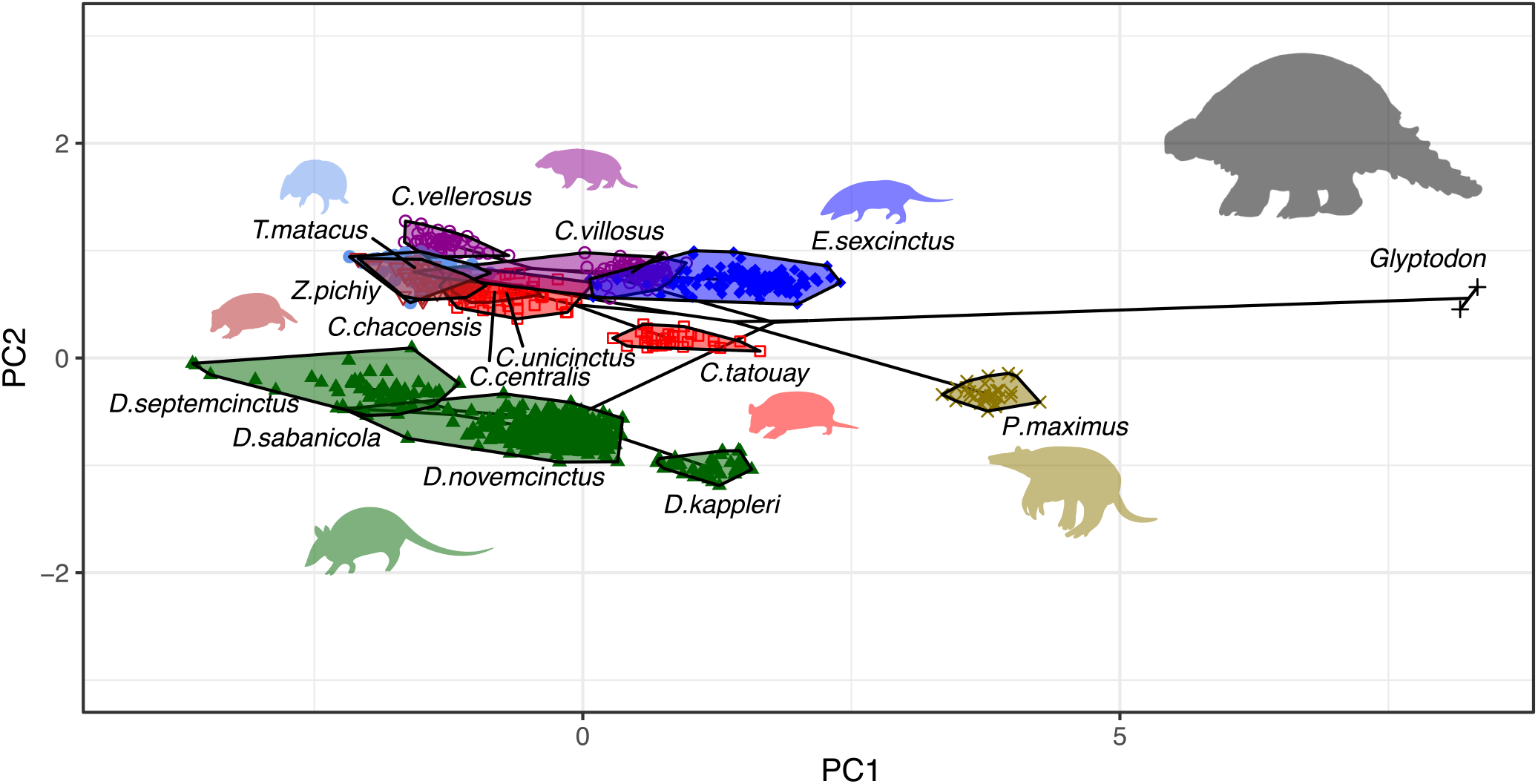
Principal component analysis (PCA) of the 33 morphometric measurements of the skull of Cingulata species. Convex polygons encapsulate each species. Symbols and colors identify genus as shown in fig. 2. Silhouettes kindly produced by M. C. Luna.

### Reconstruction of past selection

To inspect if the selective pressures necessary to generate a Glyptodont morphology differ from the remaining Cingulata in scale and direction (*i.e*., which combination of traits are favored by natural selection), we calculated the selection gradients (*β*s) necessary to generate all divergence observed in our sample (10, 17) using a version of the multivariate Breeder’s equation

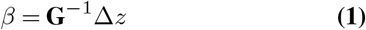

with Δ*z* being the phenotypic evolution that happened during a period of time. Here, we estimated Δ*z* as the time-standardized Phylogenetic Independent Contrasts (PIC) obtained at each node (29, 30), ensuring that *β* obtained like this are also phylogenetically independent from each other (31).

We applied an extension on eigenvalues after the 21^*st*^ to minimize the effect of noise in the calculation of *β* (32) (see supplementary material; Fig. S4). To summarize multiple *β*s, we performed an analysis similar to PCA, in which nodewise *β*s are projected on the leading lines of most selection. This was done by calculating the matrix of average cross-product of *β*s (the matrix of covariance of realized adaptive peaks), extracting the leading eigenvectors from that matrix, and projecting individual *β*s on the subspace defined by these axes (31). The number of leading vectors was evaluated employing a simulation approach (31) (see supplementary material).

### Mode of evolution

To evaluate the evolutionary processes involved in generating these morphological patterns, we employed Lande’s generalized genetic distance (LGGD), which allows confronting observed rates of evolution to the expected rate under genetic drift while taking into account the genetic association (covariances) among traits (1). The LGGD is calculated as

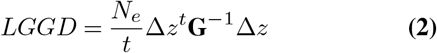

where *t* is the time in generations, and *N*_*e*_ is the effective population size. LGGDs were obtained for form, allometry-free shape features, and size.

For this analysis, the phylogeny was scaled, so the divergence time is given in generations instead of Myr. Generation times were set to be equal to the timing of sexual maturity of females plus the incubation period without assuming any reproductive seasonality. This is considered a conservative estimate, as generation times calculated like this are lower, leading to lower rates of evolution. Nevertheless, relaxing some of these assumptions (*e.g*., assuming seasonality or higher generation times) did not change our results (not shown). To obtain generation times for species without data on life-history traits (fossil species included), we estimated the linear relationship (in log scale) between body mass and generation time for 805 mammal species (33), and used the results to estimate the missing data. Because Cingulata shows a proportionally shorter generation time than the linear trend for all mammals, we subtracted a constant to correct for that difference (Fig. S2). Given that *Glyptodon* body mass estimates can vary, ranging from 819kg to 2000kg (21, 34), we calculated generation times for these extreme values. This resulted in generation times estimates ranging from 3.8-4.6 years, consistent with generation times for large ungulates (33, 35). Finally, to scale the phylogeny, we reconstructed ancestral values for generation time using maximum likelihood (36). Each branch length was then divided by the average between the ancestral and descendent generation times. PIC values obtained in the scaled phylogeny are given in units Δ*z/t*.

Since there is little available information for the *N*_*e*_ of Cingulata, we sampled *N*_*e*_ values from a uniform distribution ranging from 10,000 to 300,000. Estimates of effective population sizes for other Late Quaternary large herbivores range from 15,000-790,000 (35), so the values chosen here can be seen as conservative estimates in the case of glyptodonts. To approximate **G**, we employed the Cheverud Conjecture (37), which states that the pattern of phenotypic covariances **P** can be used as a proxy for **G** when certain conditions are met (26). Because **P**s contain more variance than **G**s, we scaled the latter according to a constant which, in the case of perfect proportionality between matrices, is equal to the average heritability 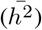 of all traits (9). Values for 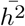 were drawn from a uniform distribution ranging from 0.3 and 0.6, which are compatible with heritability estimates for the same cranial traits in other mammalian species (38, 39). Thus, we produced a range of LGGD values for each node to account for uncertainty on both *N*_*e*_ and 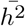.

Because parametric methods had an inadequate type I error rate (see Fig. S3), a null distribution of LGGD was constructed by simulating multivariate evolution under genetic drift as follows

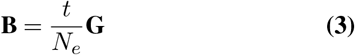

where **B** is the covariance matrix between evolutionary changes (1). For each simulation, the randomly generated value of *N*_*e*_ was used to construct a **B** from which one observation would be drawn for one generation (*t* = 1). This procedure was repeated 10000 times to generate a null distribution under drift. Values below 2.5% of the simulated values are considered to be dominated by stabilizing selection, and values above 97.5% of the simulated values are considered to be dominated by directional selection (Fig. 1). Note that estimated rates in a macroevolutionary context are better seen as the net effect of the sum of all evolutionary processes that acted during the history of a lineage (40). Thus, instead of classifying lineages under either directional or stabilizing selection (or drift), our results should be seen as an investigation of the relative importance of these multiple processes.

## Results

### Morphometrics and past selection

The principal component analysis (PCA) of our full multivariate dataset of skull measurements illustrates the uniqueness of glyptodontinae morphology (Fig. 3). Glyptodontinae shows an extreme score on PC1 compared to the extant armadillos, which is considered a size and allometric direction (Table S2); (28). The giant armadillo *Priodontes maximus* also presents a high score on PC1, but with values still closer to the remaining Cingulata than to *Glyptodon*. Dasypodidae species tend to occupy a different position of the morphospace than Euphractinae and Tolypeutinae, with similar PC1 values but smaller PC2 scores associated with an increase in the size of facial traits (Table S2).

The simulation approach designed to identify how many directions of preferential selection are necessary to produce the morphological diversity in Cingulata recovered the first three leading eigenvectors of the covariance matrix of selection gradients (*W*_1_, *W*_2_ and *W*_3_) as significant (Fig. S5). The leading vector *W*_1_ summarizes selection for the reduction of the skull’s total length while also showing a dorsoventral expansion of the face region, along with an increase in the tooth-row size (Fig. 4). The only node that scores heavily on this axis is the one associated with Glyptodonts’ origin (node 21), suggesting that this is a Glyptodont-exclusive combination of selective pressures. The second axis *W*_2_ summarizes a contrast between face and neurocranium. The nodes with the highest scores on this axis, 20 and 22, neighbor the node leading to glyptodonts, while the ones with the lowest scores, 24 and 26, are nodes internal to the *Cabassous* genus. The third axis *W*_3_ shows strong selection on features associated with the skull height (Fig. S6). An inspection of the scores along *W*_3_ reveals that this axis is mostly associated with the differentiation within Euphractinae (nodes 27—29).

**Fig. 4.**
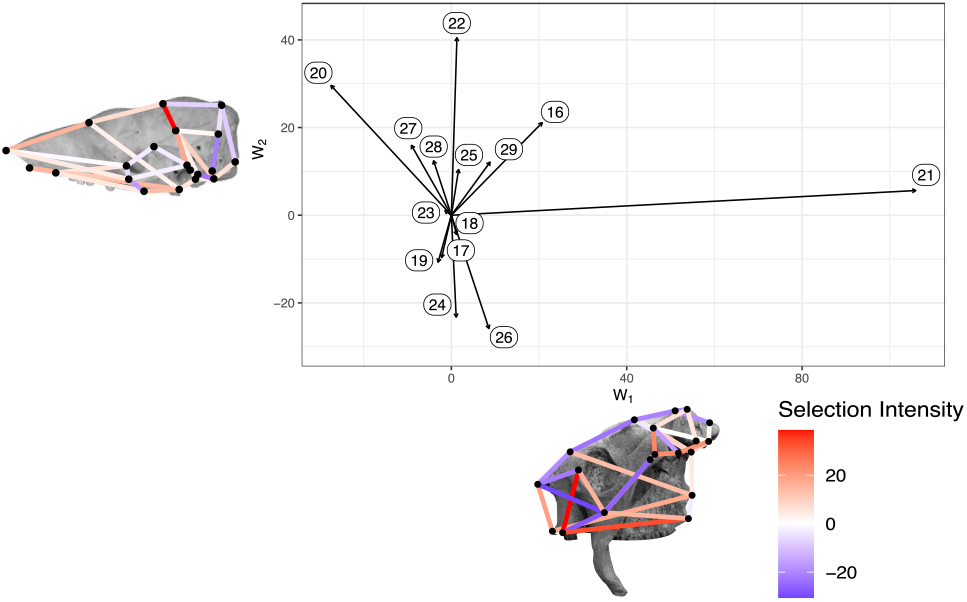
Projection of node-wise selection gradients on the two main predominant directions of directional selection (*W*_1_ and *W*_2_). Numbers represent the nodes on the phylogeny (Fig. 3A). Skulls depict a graphical representation of the two main directions *W*_1_ (*Glyptodon* skull) and *W*_2_ (*Cabassous* skull). Colors represent the intensity and direction (blue-negative and red-positive) of selection.

### Mode of evolution

LGGD shows that stabilizing selection was the dominant force in the morphological evolution of Cingulata, a fact that is more evident for both form and shape (Fig. 5A-B). For size, most observed LGGD values fell within the expected under drift (Fig. 5C). While most nodes failed to exhibit values above the expected under drift, a noticeable exception was the one representing the divergence between Glyptodontinae and Tolypeutinae (node 21). This node presented LGGD values for form, shape, and size that fall above the expected under drift, suggesting that directional selection had a preeminent role during the divergence of this group. The multivariate PIC for this node and PC1 of the full dataset (Fig. 3) are highly aligned (vector correlation=*>* 0.98), indicating that this direction is mostly associated with the divergence of Glyptodonts from all remaining Cingulata. The increased evolutionary rate observed for this node does not seem to be an artifact introduced by the timing of the origin of Glyptodonts, as older nodes associated with the divergence of subfamilies within Cingulata did not show elevated LGGD values (Fig. 5).

**Fig. 5.**
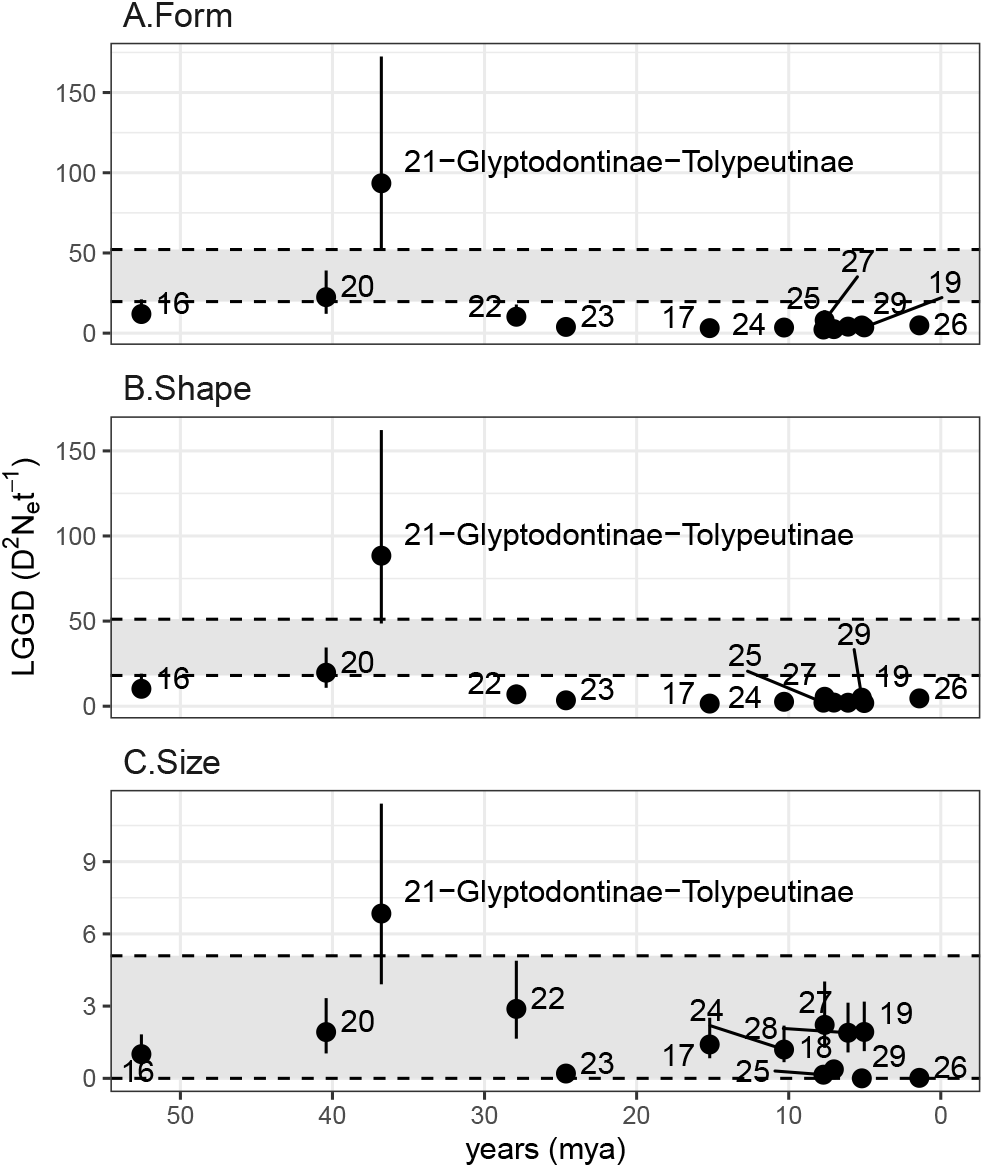
Values for Lande’s Generalized Genetic Distance (LGGD) calculated at each node plotted against node age for form (**A**.), shape (**B**.) and size (**C**.). Dots and solid lines represent the median and the ranges for the LGGD values for a given node. Nodes are numbered following fig. 3A. Gray area represents the expectation under drift. Values above it are consistent with directional selection and below it with stabilizing selection.

## Discussion

Previous investigations on the macroevolutionary rates of morphological evolution in mammals have shown that morphological change is usually orders of magnitude slower than would be expected by drift (5). Such macroevolutionary rates are usually measured as the ratio between total morphological differentiation and time (29). As divergence times increase, even extreme changes can get diluted, implying that searching for signals of directional selection in macroevolutionary data might be futile (Fig. 1). In this sense, or analysis produce a more nuanced understanding of such process.

Broadly speaking, our results agree with previous evidence that stabilizing selection is the dominant evolutionary process shaping mammalian morphological diversification (5, 7, 9). Rates of morphological evolution calculated throughout the evolutionary history of Cingulata are consistent with stabilizing selection, a pattern that is more expressive for shape than for size (4). This finding is in line with previous suggestions that shape features are more constrained than size, implying that the former is more affected by stabilizing selection than the latter (41). One proposed explanation for this pattern refers to the macroevolutionary dynamic of adaptive zones (42, 43). Adaptive zones are regions of the phenotypic space defined by a specific ecomorphological association between phenotype, function and ecology. Within that region, evolution normally results in quantitative differences in ecology and morphology. When lineages escape such region, thus invading a new adaptive zone, this is usually associated with more intense changes in morphology and ecology, a feature usually observed in major evolutionary transitions and inovations (42, 43). If, as some have suggested, shape features are more strongly associated with functional demands (44–48), than changes in functional requirements would probably lead to shape evolution, and possibly to the invasion of a new adaptive zone (31, 47). Size changes, on the other hand are less likely to impact ecomorphological relationships, being associated with within-zones evolution (31, 47). In fact, most of the evolution of Cingulata seems to concentrate on a similar region of the morphospace (Fig. 3), which could be evidence that their morphology is being maintained within a “generalist-armadillo” adaptive zone (45). Within that zone, shape and size would be under strong stabilizing selection, but size would be more prone to change

The only exception to this rule, both in terms of morphospace occupation and rates of evolution, is the glyptodont lineage (Fig. 3, 5). Glyptodonts seem to have reached a region unoccupied by other Cingulata (Fig. 3). The most straightforward explanation for this is that glyptodonts invaded a new adaptive zone fuelled by a drastic change in ecology. Glyptotonts are thought to be highly specialized herbivores, a diet with strong functional demands (21). These demands were then responsible for imposing a strong directional selection, leading to radical morphological change and functional innovation seen in this group (20). This interpretation is further reinforced by the investigation of hypothetical past selection. The description of *W*_1_, which is a Glyptodont-exclusive axis, matches the telescoping process that Glyptodonts went through during their evolutionary history (20). Furthermore, the required strength of selection for the divergence of glyptodonts is superior to those observed within the generalist-armadillo adaptive zone. Changes in the orientation and strength of selection have been associated with dietary shifts in other mammals (31, 49), reinforcing this idea that the selective change observed in Glyptodonts has an ecomorphological basis.

The fact that the signal of directional selection is detectable in a macroevolutionary scale is remarkable. To our knowledge, a comparable signal was only previously found at the divergence between lower and higher apes, at 20mya, and at the origin of humans, at 6mya (9). This is somewhat expected, as primates are marked by a phylogenetic trend towards increased brain size, a pattern that is intensified on hominins (50). It is possible that the evolution of Glyptodontinae was also marked by a continuous differentiation trend. Early representatives of the group, such as *Propalaehoplophorus*, exhibit an intermediary version of the cranial telescoping characteristic of latter glyptodonts (20). Given that the split between Glyptodontinae and Tolypeutinae happened 35mya ago (24, 25) and *Propalaehoplophorus* is known from the fossil record from 22-15mya (51), this suggests that the accumulation of morphological changes in this lineage was probably gradual and continuous. If that is the case, then glyptodonts can be the oldest and longest case of such a pattern of continuous change. Alternatively, if the group’s evolution was punctuated by long periods of stasis, this could mean that directional selection events were even more intense than shown by the present analysis (Fig. 1). However, given the clear difference between primitive and derived forms within the group, the truth probably lies somewhere in between, with a gradual directional evolution punctuated by some periods of stasis but in proportions that differ from the remaining Cingulata and even other mammals.

As mentioned above, the use of biologically informed models to investigate evolutionary rates at macroevolutionary scales is still rare and limited in scope. We suggest that some of this dismissal comes from the assumed inapplicability of these models at larger time scales due to the overwriting power of stabilizing selection. Here we show that this assumption does not hold for glyptodonts, raising the possibility that directional selection is more pervasive than previously thought. The integration of more refined paleontological, demographic, and life-history data in the investigation of macroevolutionary questions can lead to a more nuanced view of the role of directional and stabilizing selection on the evolution of complex morphologies.

## ACKNOWLEDGEMENTS

We thank the curators of the following institutions for providing access to specimens and support for data acquisition: American Museum of Natural History; California Academy of Science; Faculdad de Ciencias de la Universidad de la Republica; Field Museum of Natural History; Museu Paraense Emílio Goeldi; Idaho Museum of Natural History; Los Angeles County Museum; La Brea Tar Pits and Museum; Museo Argentino de Ciencias Naturales; Museu de Ciências Naturais da Pontifícia Universidade Católica de Minas Gerais; Museu de Ciências Naturais da Fundação Zoobotânica do Rio Grande do Sul; Museo de La Plata; Museo Paleontológico de Colonia del Sacramento; Museu Nacional de Historia Natural de Montevideo; Museu Nacional (Rio de Janeiro); University of California Museum of Vertebrate Zoology; Natural History Museum (London); University of California Museum of Paleontology; Florida Museum of Natural History; Smithsonian National Museum of Natural History; Museu de Zoologia da Universidade de São Paulo; Universidade Federal do Piauí. We also thank EvoClub_BR discussion group and the members of Uyeda’s Lab (Virginia Tech) for thoughtful comments and suggestions to the text. This work was partially supported by grants from NSF to FAM (DEB 1350474 and DEB 1942717) and FAPESP to AH and GM (2012/24937-9; 2011/14295-7).

## Supplementary methods and Results

### Sample and measurements

We obtained morphometric variables from skulls belonging to 14 extant and one extinct species, representing four Cingulata subfamilies (Table S1). Only adult individuals were sampled. Criteria used to determine adulthood are explained in (26).

Landmarks are the same as in (26, Fig. S1). The set o measurements refer to the 35-measurements found in (26) minus two measurements: PT-TSP and APET-TS. These measurements were excluded because they can sometimes reach small values on some species, and their variance is disproportionately inflated on the log scale. See (26) for more information on landmark and measurement description, data processing, and measurement repeatabilities.

### Phylogeny

The phylogenetic relationships between species were based on molecular analysis on mitogenomic data from Delsuc et al. (24; Fig. 3C), consistent with other independently derived analyses (25). Both papers determined the placement of Gliptodonts on the Xenarthran phylogeny based on ancient DNA samples extracted from *Doedicurus* sp specimens. Here, we swapped *Doedicurus* sp. for *Gliptodon* sp. because there is ample evidence showing that both belong to the same subfamily (19, 22). In the absence of any other member of the group in our analysis, the phylogenetic position of one glyptodontid can be assigned to any other species of that clade. Furthermore, because both species are thought to have gone extinct at roughly the same time (around the end of the Pleistocene and the beginning of the Holocene), there is no need to scale branch lengths accordingly.

### Generation time

Generation times were obtained (33) by summing up the timing of sexual maturity of females and the incubation period. Because armadillos reproduce more than once throughout their lifetimes, real generation times are probably larger than this. As a consequence of our calculations, the values used here for generation times are considered a minimum benchmark value and not the true one. Furthermore, because larger generation times imply that fewer generations have transpired during the same period, this means that our estimates of LGGD are probably underestimated. Given that small LGGD are consistent with stabilizing selection and that stabilizing selection is expected to be the dominant force in the evolution of morphological traits, the use of these underestimated generation times leads to conservative estimates of LGGDs.

**Table S1.** Sample size by species.

| Subfamily | Species | sample |
| --- | --- | --- |
| Dasypodinae | *Dasypus kappleri* | 34 |
|  | *Dasypus novemcinctus* | 341 |
|  | *Dasypus sabanicola* | 2 |
|  | *Dasypus septemcinctus* | 52 |
| Tolypeutinae | *Cabassous centralis* | 9 |
|  | *Cabassous chacoensis* | 2 |
|  | *Cabassous tatouay* | 30 |
|  | *Cabassous unicinctus* | 37 |
|  | *Tolypeutes matacus* | 39 |
|  | *Priodontes maximus* | 30 |
| Glyptodontinae | *Glyptodon* | 2 |
| Euphractinae | *Chaetophractus vellerosus* | 47 |
|  | *Chaetophractus villosus* | 65 |
|  | *Euphractus sexcinctus* | 125 |
|  | *Zaedyus pichiy* | 45 |
|  | **TOTAL** | **860** |

**Fig. S1.**
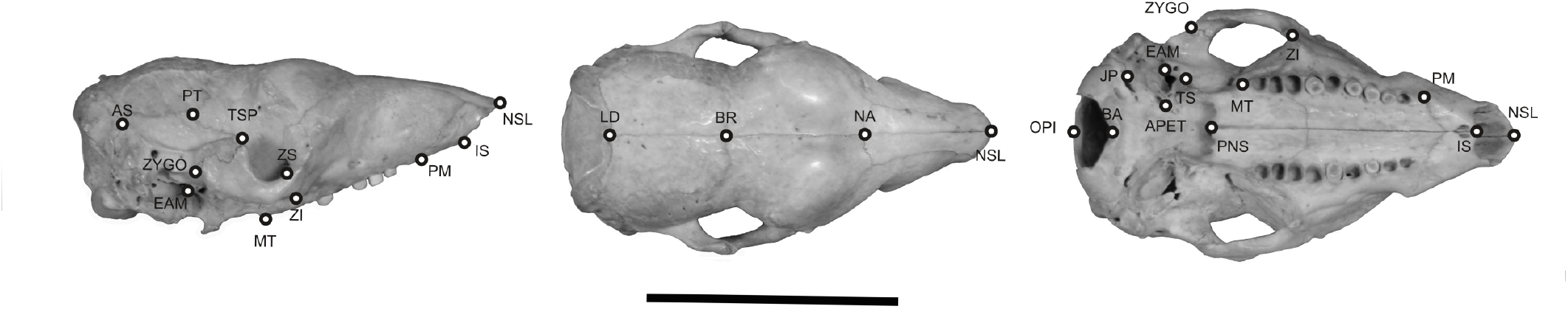
Cranial landmarks measured in this study represented on a *Cabassous* specimen. Bar is equal to 5 cm.

**Table S2.** Loadings of the first two principal components of the intra and interspecific morphometric datasets. Traits are discriminated according to their cranial anatomical regions.

| Trait | Anatomical region | Intraspecific |  | Interspecific |  |
| --- | --- | --- | --- | --- | --- |
|  |  | PC1 | PC2 | PC1 | PC2 |
| IS.PM | Face | 0.194 | 0.058 | 0.178 | -0.017 |
| IS.NSL | Face | 0.200 | 0.012 | 0.200 | -0.031 |
| IS.PNS | Face | 0.175 | 0.033 | 0.175 | -0.209 |
| PM.ZS | Face | 0.169 | 0.159 | 0.153 | -0.337 |
| PM.ZI | Face | 0.165 | -0.041 | 0.139 | -0.315 |
| PM.MT | Face | 0.162 | 0.054 | 0.172 | -0.114 |
| NSL.NA | Face | 0.183 | 0.101 | 0.178 | -0.197 |
| NSL.ZS | Face | 0.177 | 0.132 | 0.160 | -0.225 |
| NSL.ZI | Face | 0.175 | -0.016 | 0.153 | -0.217 |
| NA.PNS | Face | 0.170 | 0.002 | 0.175 | -0.123 |
| PT.ZYGO | Face | 0.219 | 0.000 | 0.195 | -0.118 |
| ZS.ZI | Face | 0.213 | -0.904 | 0.126 | -0.029 |
| ZI.MT | Face | 0.212 | -0.009 | 0.189 | 0.275 |
| ZI.ZYGO | Face | 0.197 | 0.291 | 0.239 | 0.298 |
| ZI.TSP | Face | 0.244 | 0.138 | 0.235 | 0.207 |
| MT.PNS | Face | 0.181 | -0.055 | 0.190 | -0.340 |
| EAM.ZYGO | Face | 0.181 | -0.027 | 0.195 | 0.095 |
| ZYGO.TSP | Face | 0.236 | 0.004 | 0.213 | 0.166 |
| NA.BR | Neurocranium | 0.171 | 0.021 | 0.161 | -0.162 |
| BR.PT | Neurocranium | 0.090 | -0.009 | 0.126 | -0.021 |
| BR.APET | Neurocranium | 0.141 | 0.012 | 0.142 | 0.019 |
| PT.APET | Neurocranium | 0.136 | 0.015 | 0.141 | 0.041 |
| PT.BA | Neurocranium | 0.140 | 0.005 | 0.157 | 0.048 |
| PT.EAM | Neurocranium | 0.160 | -0.003 | 0.181 | 0.076 |
| PNS.APET | Neurocranium | 0.204 | 0.075 | 0.189 | 0.015 |
| APET.BA | Neurocranium | 0.139 | 0.009 | 0.174 | 0.131 |
| BA.EAM | Neurocranium | 0.140 | 0.027 | 0.180 | 0.287 |
| LD.AS | Neurocranium | 0.139 | -0.013 | 0.192 | 0.174 |
| BR.LD | Neurocranium | 0.133 | -0.033 | 0.151 | 0.106 |
| OPI.LD | Neurocranium | 0.177 | 0.044 | 0.144 | -0.026 |
| PT.AS | Neurocranium | 0.160 | -0.024 | 0.200 | 0.141 |
| JP.AS | Neurocranium | 0.161 | 0.037 | 0.146 | 0.043 |
| BA.OPI | Neurocranium | 0.076 | -0.010 | 0.115 | -0.059 |

### Type I error rates for LGGD

To access the adequacy of the parametric test proposed for Lande’s Generalized Genetic Distance (1), we employed a simulation approach. We generated 10,000 rounds of one generation neutral multivariate evolution using equation 3 and known parameters for 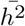 and *N*_*e*_. For each simulated vector of divergence, we resampled **P** using a MonteCarlo approach to simulate sampling error. **P**s were reestimated with the same original empirical sample (n=860). Because the calculation of LGGDs involves the inversion of the **G**, we also evaluated if a matrix extension could improve the estimation of LGGDs. The simulated and observed values of LGGD were confronted against the 95% interval under the null hypothesis following a *χ*-square distribution with 33 degrees of freedom (1).

Results show that in the original simulated LGGD (calculated directly from the simulated divergence and input parameters), the rejection rate of the null hypothesis using the parametric method is close to the nominal rate (rate=0.053). However, with sampling error, larger LGGDs seem to be moderately exaggerated (Fig. S3A), leading to inflated type I error rates (rate of rejection of the null hypothesis=0.062). Extensions of the **G** prior to inversion led to underestimated LGGDs (Fig. S3B) and to even higher type I error rates (rates=0.669). This result shows that matrix extension does not constitute a solution to sampling errors in this case (but see below). Therefore, we adapted a simulation approach to construct a non-parametric confidence interval for the null hypothesis of evolution under drift.

### Matrix inversion and extension

To evaluate the potential effects of sampling error of **G** on the estimation of selection gradients (*β*), we performed two analyses. The first is the evaluation of the noise-floor limit, as suggested by (32). In this analysis, we evaluate the behavior of the second derivative of a discrete function defined as the relationship between the rank of an eigenvalue and the variance of eigenvalues defined by binning five neighboring eigenvalues. The noise floor limit is reached when the second derivative of this function approaches zero, and the eigenvalue rank in which this occurs is set to be the cutoff value for the extension method (32). Here we defined the noise-floor region at 1*e* 07, suggesting that the first 21 eigenvalues should be maintained, and the remaining should be extended (Fig. S4A).

**Fig. S2.**
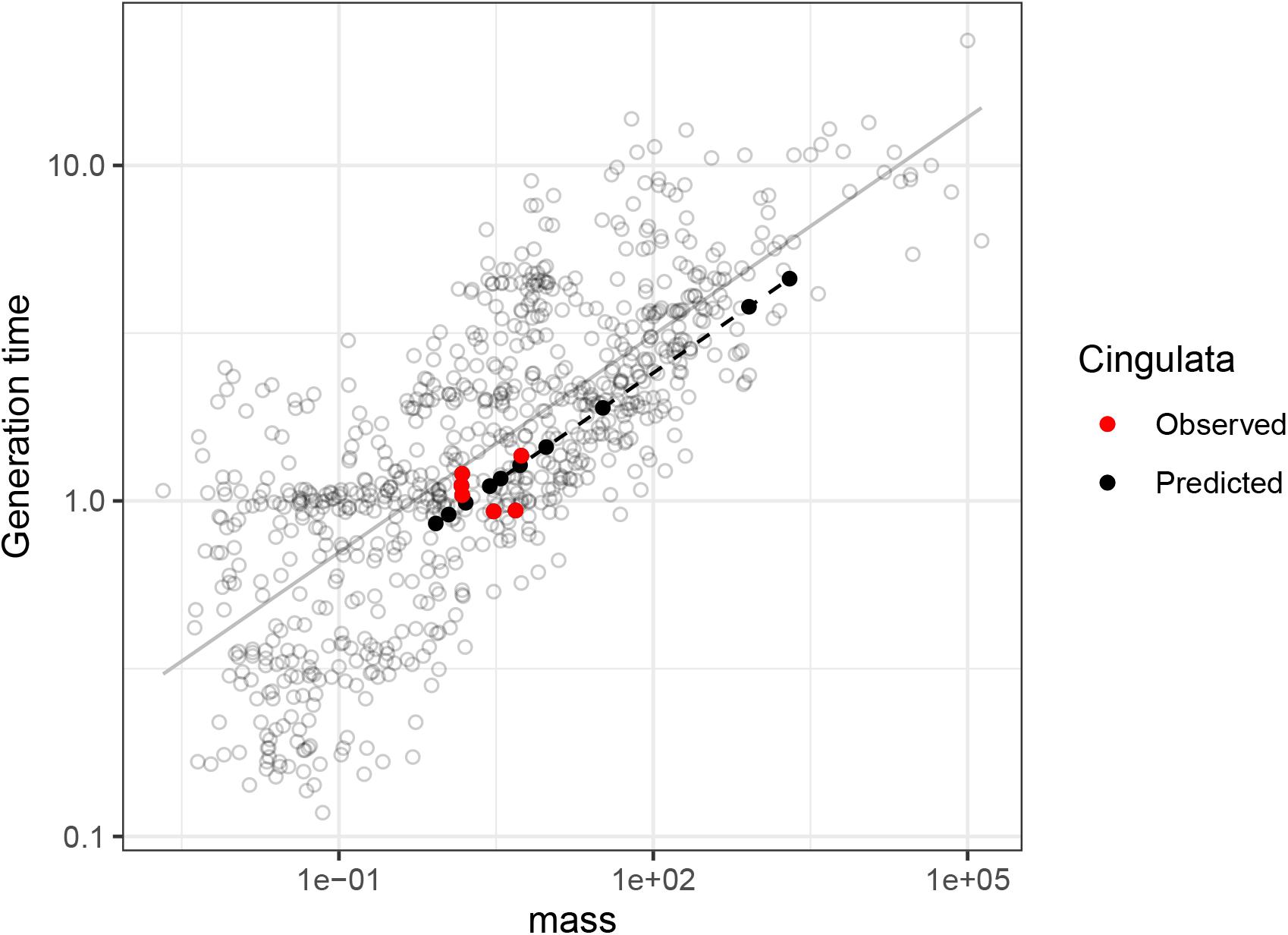
Relationship between generation time and body mass for 805 mammalian species. The solid gray line represents the linear relationship for all species. Red-filled circles represent Cingulata taxa with both mass and generation time information. Black-filled circles represent Cingulata taxa with only mass and with a predicted generation time. The dashed line represents the model adopted for the association between mass and generation time among Cingulata.

**Fig. S3.**
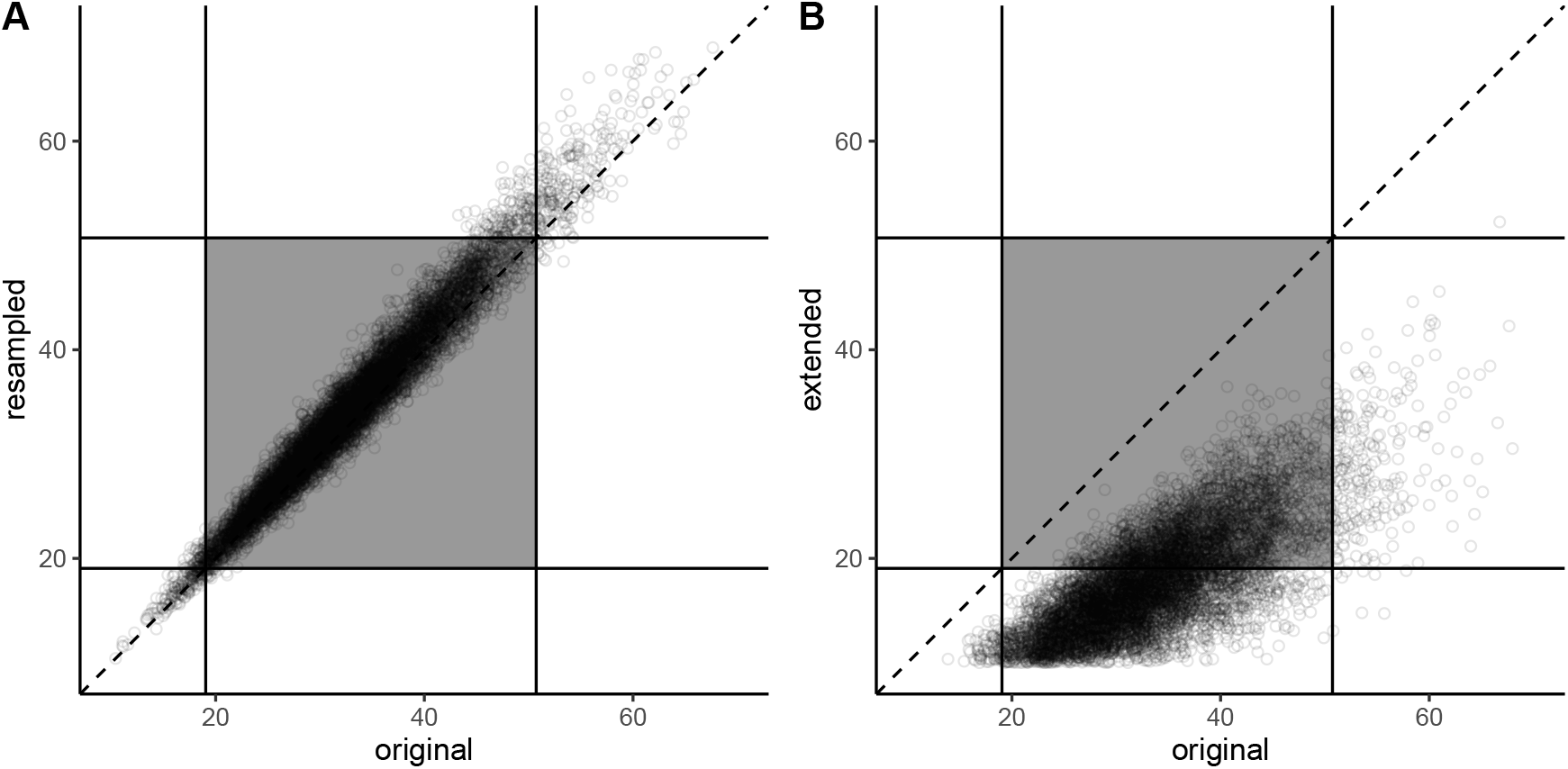
Relationship between actual (original) and estimated LGGD for resampled matrices without (**A**) and with extension (**B**), respectively. The dark quadrant represents the 95% theoretical intervals for the null-hypothesis under drift for both the original and resampled matrices. The dashed line represents the line in which observed and original values are equal.

**Fig. S4.**
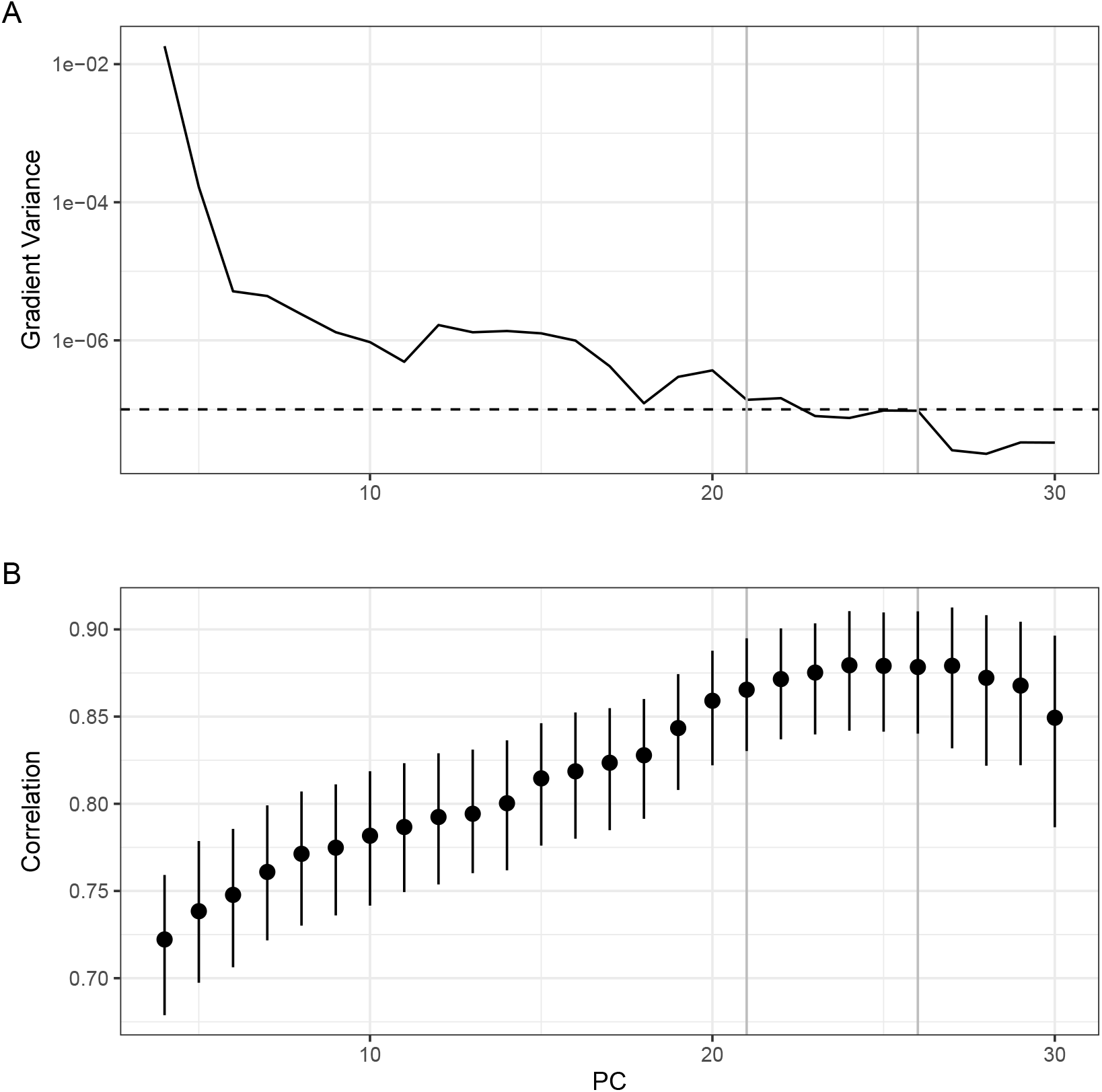
**A**. Noise floor analysis for the pooled within-group covariance matrix for Cingulata. **B**. Correlation between original and estimated *β*s on the simulation approach. Horizontal grey lines represent numbers of eigenvalues retained that show high vector correlation between original and estimated *β*s and are near the noise floor.

To validate this analysis, we replicated (32) simulations, in which a known *β* was re-estimated on a new matrix resampled adopting a Monte Carlo approach (see 31 as well). We performed this simulation by changing the number of retained eigen-values from 6 to 30. In each case, we performed the simulation 1000 times (25,000 simulations in total). For each simulation, we calculated the vector correlation between the original and the estimated *β*. Higher values imply that the estimated *β* is in the same direction as the original simulated selection gradient. From 6 to 21 eigenvectors, the correlation between original and estimated selection gradients increased almost steadily, reaching a plateau from 21 to 26 eigenvalues (Fig. S4B). After 26 vectors, the vector correlation decreases, suggesting that retaining more vectors leads to badly estimated *β*s. We chose to keep 21 vectors because it is the first eigenvalue to reach the noise-floor on the previous analysis. Furthermore, estimations of the selection gradient to generate a glyptodont morphology did not significantly change with the retention of more axes.

### Adaptive landscape and selective lines of least resistance

In a quantitative genetics framework, adaptive landscapes can be investigated as the variance-covariance matrix of adaptive peaks (17, 30, 31). Following (31), we calculated this matrix as follows

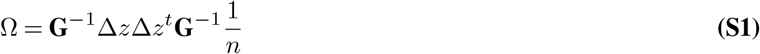

**Fig. S5.**
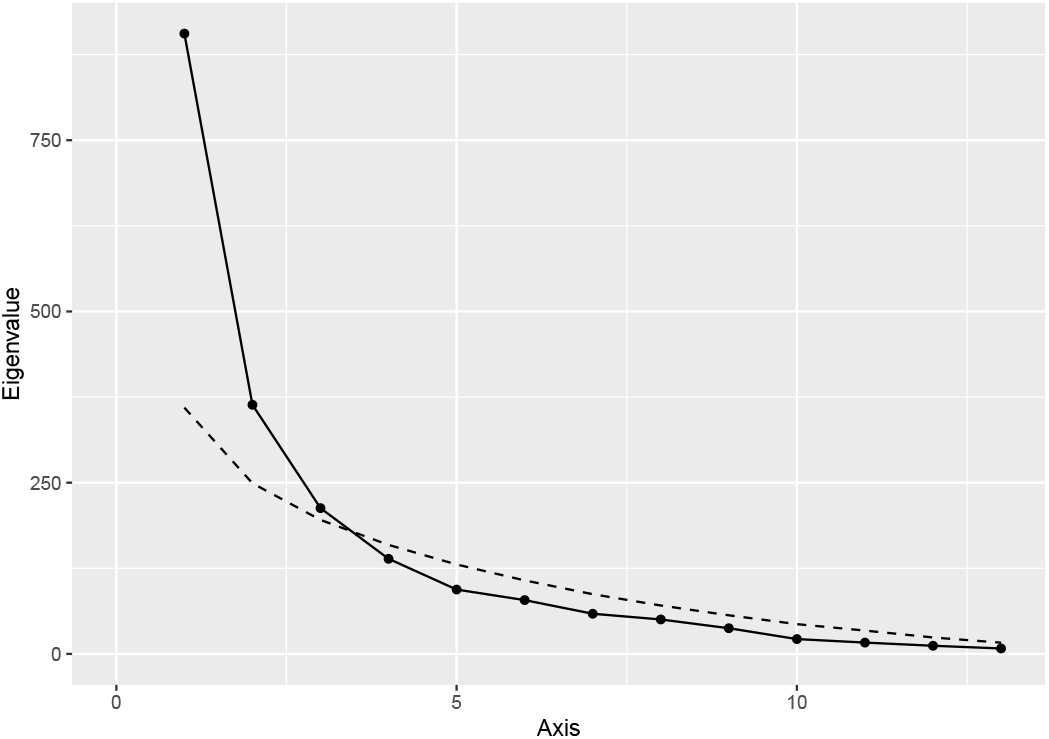
Empirical (solid line) and simulated (dotted line) distribution of Ω’s eigenvalues.

With n being the species sample size. This equation is equal to the cross-products of the selection gradients. The *ith* eigenvectors of Ω (*W*_*i*_) can be interpreted as predominant directions of directional selection, and the projection of node-specific *β*s can be interpreted as the loading of a specific gradient to that axis.

Following (31), we performed a Monte-Carlo simulation approach to investigate which axes could be considered different from neutral evolution. This was done using equation 3to generate 10,000 datasets under drift. For each round of the simulation, Ω was calculated and its eigenvalues were obtained. The resulting Ω was scaled to have the same trace as the empirical one, so we only consider the distribution of eigenvalues, not their absolute magnitude. The distribution of eigenvalues of the simulations was then confronted to the empirical eigenvalues, and those greater than 95% than the simulated ones were considered significant.

Because the Glyptodont divergence dominates the dataset, we performed two different sets of simulations. One where we included Glyptodont to calculate the empirical trace of Ω, and one where it was excluded. For the simulations including Glyptodont, only the first leading eigenvalue was considered significant, while the exclusion of glyptodonts showed that the three leading axes are significant (Fig. S5). An investigation of the scores of node-specific *β*s on *W*_3_ shows that the only nodes with high scores are nodes 28 and 29 (Fig. S6A). These nodes represent the divergences among *Chaetophractus* and *Zaedyus* species. The multivariate representation of the selection gradients shows mainly a concentrated pressure to expand the nasal cavity and increase the distance between the anterior-most portion of the zygomatic arch and the tip of the snout (Fig. S6B). This representation matches the described differences of *Zaedyus* in relation to *Chaetophractus*. Because this axis is so phylogenetically restricted, we focus on the leading two directions on the main text.

**Fig. S6.**
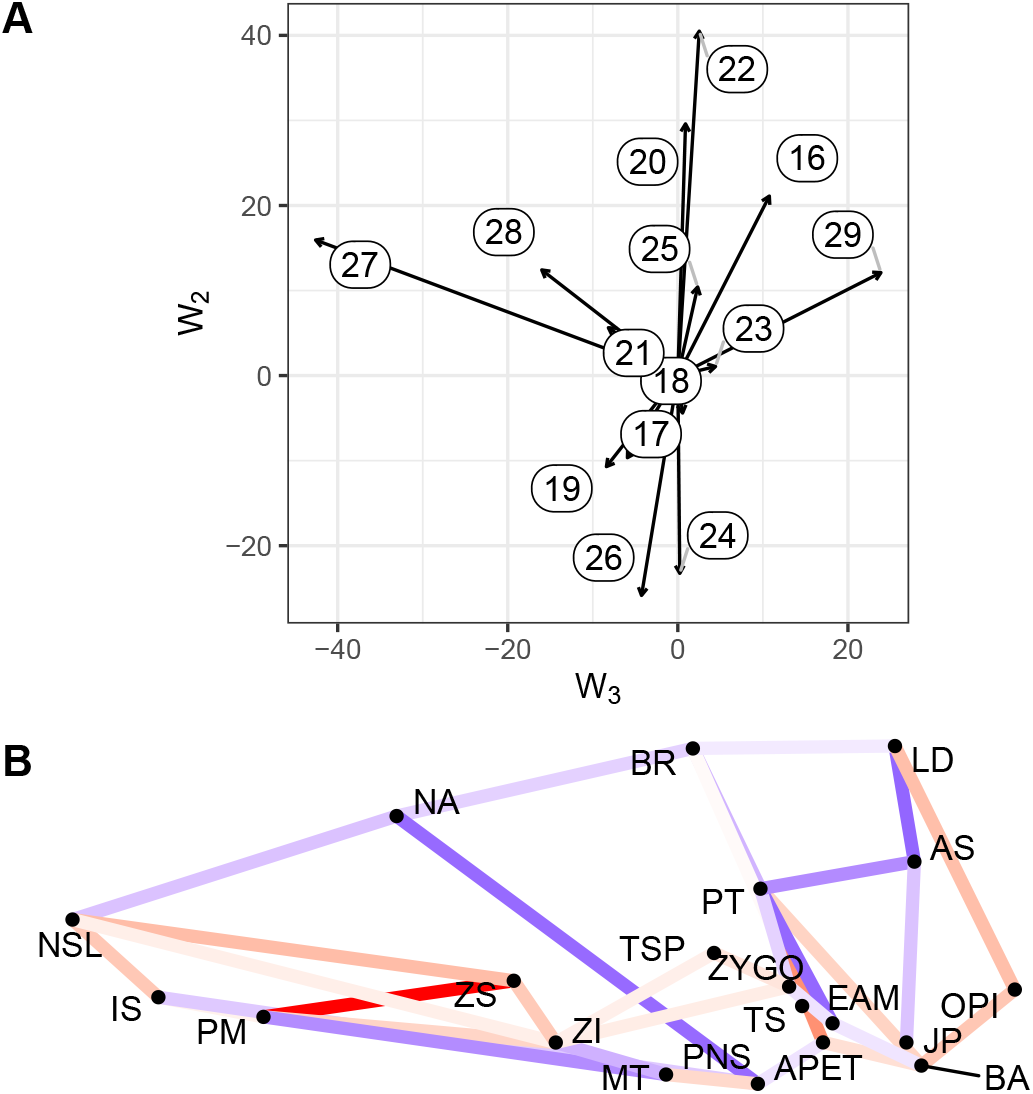
**A**. Projection of node-wise selection gradients on the two main predominant directions of directional selection (*W*_3_ and *W*_2_). Numbers represent the nodes on the phylogeny (Fig. 3A). **B**. Graphical representation of *W*_3_. Color represents the intensity of positive (red) and negative (blue) selection, as in Fig. 4.

